# *apterous A* specifies dorsal wing patterns and sexual traits in butterflies

**DOI:** 10.1101/131011

**Authors:** Anupama Prakash, Antónia Monteiro

## Abstract

Butterflies have evolved different color patterns on their dorsal and ventral wing surfaces to serve different signaling functions, yet the developmental mechanisms controlling surface-specific patterning are still unknown. Here, we mutate both copies of the transcription factor *apterous* in *Bicyclus anynana* butterflies using CRISPR/Cas9 and show that *apterous A* functions both as a repressor and modifier of ventral wing color patterns, as well as a promoter of dorsal sexual ornaments in males. We propose that the surface-specific diversification of wing patterns in butterflies proceeded via the co-option of *apterous A* into various gene regulatory networks involved in the differentiation of discrete wing traits. Further, interactions between *apterous* and sex-specific factors such as *doublesex* may have contributed to the origin of sexually dimorphic surface-specific patterns. Finally, we discuss the evolution of eyespot pattern diversity in the family Nymphalidae within the context of developmental constraints due to *apterous* regulation.

**Significance statement:** Butterflies have evolved different wing patterns on their dorsal and ventral wing surfaces that serve different signaling functions. We identify the transcription factor, *apterous A*, as a key regulator of this surface-specific differentiation in butterflies. We also show a role for *apterous A* in restricting the developmental origin of a novel trait, eyespots, to just the ventral wing surface. Dorsal-ventral differentiation of tissues is not just restricted to butterfly wings but occurs in many other organs and organisms from arthropods to humans. Thus, we believe that our work will be of interest to a diverse group of biologists and layman alike interested in the role of development in shaping biodiversity.

## Main Text

Butterflies are a group of organisms well known for their diverse and colorful wing patterns. Due to the dual role these patterns play in survival and mate selection, many butterflies have evolved a signal partitioning strategy where color patterns appearing on the hidden dorsal surfaces generally function in sexual signaling, whereas patterns on the exposed ventral surfaces most commonly serve to ward off predators (1, 2) (Fig 1A). While the molecular and developmental basis of individual pattern element differentiation, such as eyespots or transverse bands, has been previously studied (3, 4), the molecular basis of dorsal and ventral surface-specific color pattern development remains unknown. Elucidating this process will help us understand the mechanism of diversification and specialization of wing patterns within the butterfly lineage.

**Figure 1:**
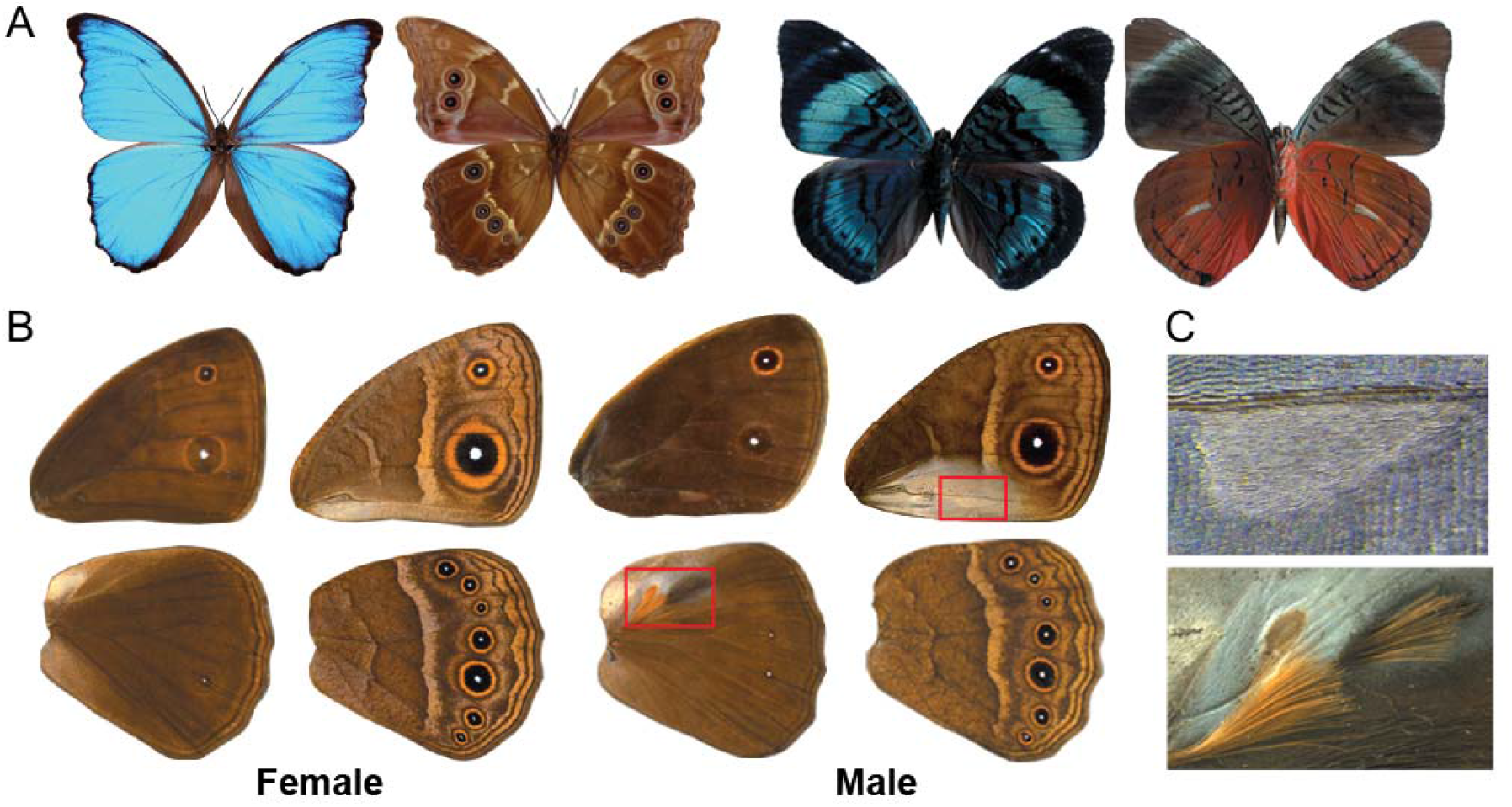
Dorsal-Ventral surface-specific variation in butterflies. A) Dorsal (left) and ventral (right) surfaces of *Morpho menelaus* and *Panacea regina* illustrating striking variation in color and patterns between surfaces. B) Dorsal (left) and ventral (right) surfaces of a male and female *Bicyclus anynana*. The regions boxed in red are expanded in C. C) Magnified view of the androconial organs present only in males. Top: Forewing ventral androconia with a characteristic teardrop shape surrounded by silver scales. The scales on the corresponding dorsal forewing surface are completely brown. Bottom: Hindwing dorsal androconia, also surrounded by silver scales, along with two patches of hair-pencils. These traits are absent from the ventral hindwing.

We hypothesized that the transcription factor *apterous (ap)*, a gene expressed on the dorsal wing surfaces of flies (5), might be implicated in differentiating dorsal from ventral wing patterns in butterflies. In insects, however, this gene is often present in two copies, *apA* and *apB*, that don’t necessarily share the same expression patterns, and flies are unusual for having lost one of these copies. In the beetle *Tribolium castaneum*, *apA* is expressed on the dorsal surface whereas *apB* is expressed on both surfaces (6). In the butterfly *Junonia coenia, apA* is expressed on the dorsal surface of larval wings (7) but, the expression of *apB* and the role of either *apA* or *apB* in wing development and patterning is not known for this or any butterfly species.

## Results

### *apA* and *apB* are both expressed on dorsal surfaces of developing wings

To investigate *ap* expression in butterflies, we cloned both *ap* homologs from the African squinting bush brown *Bicyclus anynana* (Fig 1B, C), and used *in situ* hybridization to localize *apA* and *apB* mRNA in developing larval and pupal wing discs. Both homologs of a*p* were localized to the dorsal surfaces of the wings (Fig 2D, S1B). In the last larval instar wing discs, *apA* was expressed uniformly on the wing surface but absent in future dorsal eyespot centers of hindwings (Fig 2A) and forewings (Fig 2B). In larval wing discs of the *B. anynana* “Spotty” mutant, which develops two additional dorsal eyespots, *apA* was absent in the additional centers (Fig 2B). Furthermore, pupal wing expression of both *apA* and *apB* was up-regulated in dorsal male-specific cells that give rise to long and thin modified scales, the hair-pencils, used for dispersing pheromones during courtship (Fig 2C, S1C). This pattern of expression was not seen in developing female pupal wings, which lack hair-pencils (Fig 2C, S1C). Control sense probes for both *apA* and *apB* (Fig S1) did not show any surface-specific or hair-pencil specific staining patterns.

**Figure 2:**
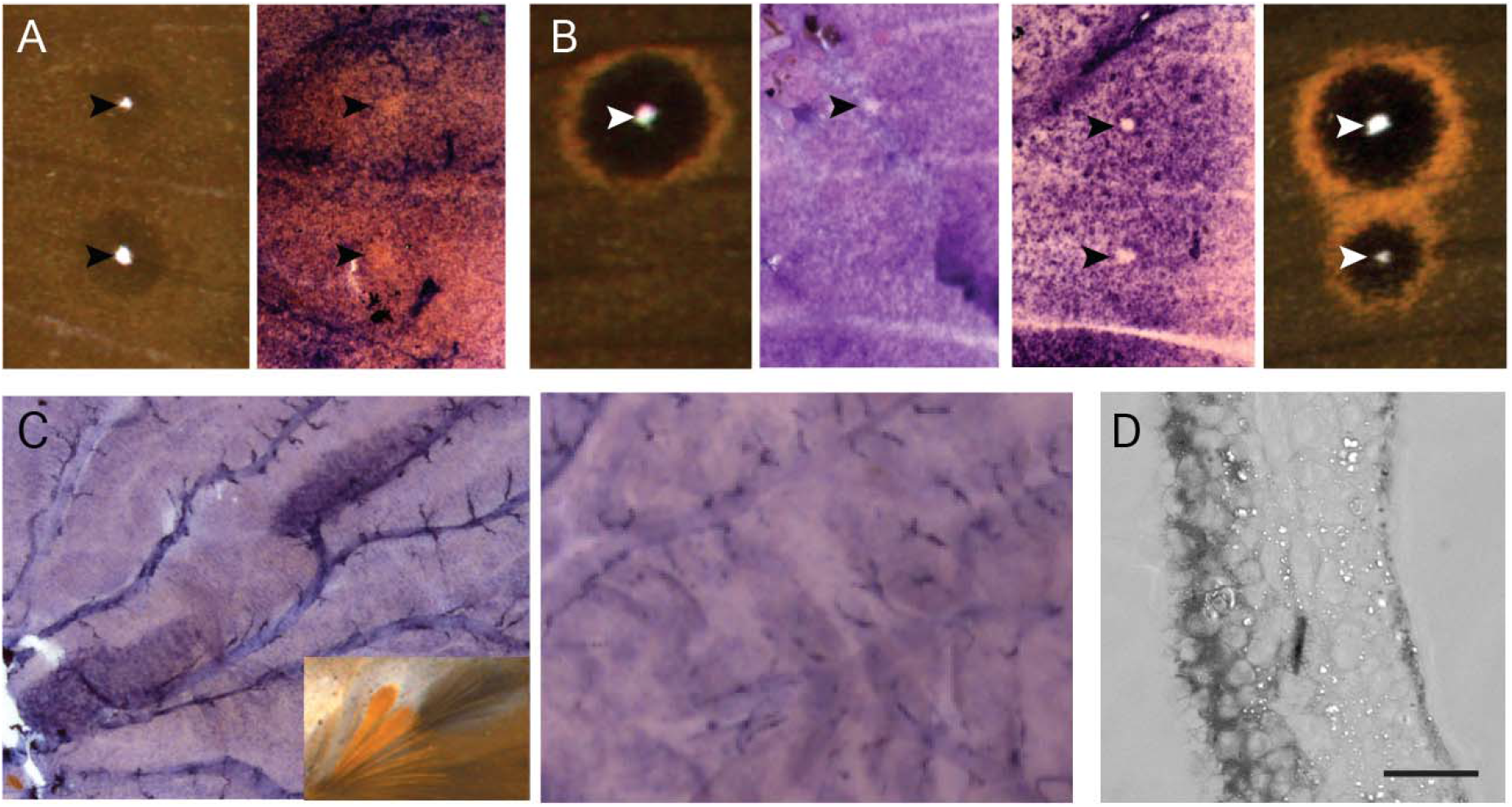
*apA* mRNA localization in developing wing discs of *Bicyclus anynana*. A) *apA* expression is uniform across the epidermis but absent in future dorsal eyespot centers of hindwings. B) *apA* expression is absent in the future dorsal eyespot center of the wildtype forewing (left) and also in the additional eyespot center in the *B. anynana* “Spotty” mutant (right). C) Male wings (left) (28 hours after pupation) showing up-regulated dorsal *apA* expression in the hair-pencil regions. Inset shows the hair-pencils in adult male *B. anynana*. Female wings (right) (25 hours after pupation) show no up-regulation of *apA* in corresponding regions of the dorsal surface. D) Cross-sectional view of a developing wing disc showing dorsal-specific *apA* expression (left side of the cross-section). Scale bar is 20µm.

### *apA* regulates dorsal surface-specific wing patterning

To functionally test the role of *ap*, we used the CRISPR/Cas9 system to disrupt the homeodomain and LIM domain of *apA* (Fig 3A) and the LIM domain of *apB* (Fig S2A) (Table S2). A range of mosaic phenotypes were observed in both types of *apA* mutant individuals (Fig 3). A few of these lacked wings, whose absence was visible upon pupation (Fig S3: mutant from batch#9, individual #1(M9-1)), and some adults had mosaic patches of ventral-like scales appearing on the dorsal surface (Fig 3B:M9-2). In other mutants, the sex pheromone producing organ, the androconial organ, of the ventral forewing appeared on the dorsal surface in males with its associated silver scales (Fig 3B:M9-27). Males also had modified hair-pencils associated with the dorsal androconial organ of the hindwing, with loss of characteristic ultrastructure and coloration, and absence of surrounding silver scales (Fig 3B:M9-12 (bottom)). Extreme mutant individuals showed improper wing hinge formation, entire wing dorsal to ventral transformation (Fig 3B: M9-3), the appearance of the ventral white band on the dorsal surface (Fig 3B:M9-12 (top)), and in one case, all seven eyespots on the dorsal hindwing (Fig 3B:M9-12 (bottom)), a surface that normally exhibits, on average, zero to one eyespot in males and one to two eyespots in females. *apA* clones also led to an enlarged outer perimeter to the gold ring in dorsal hindwing and forewing eyespots (Fig 3B:M235-11). CRISPR/Cas9 disruption effects on the target sequence were verified in a few individuals, which showed the presence of deletions in the targeted regions (Fig 3A).

**Figure 3:**
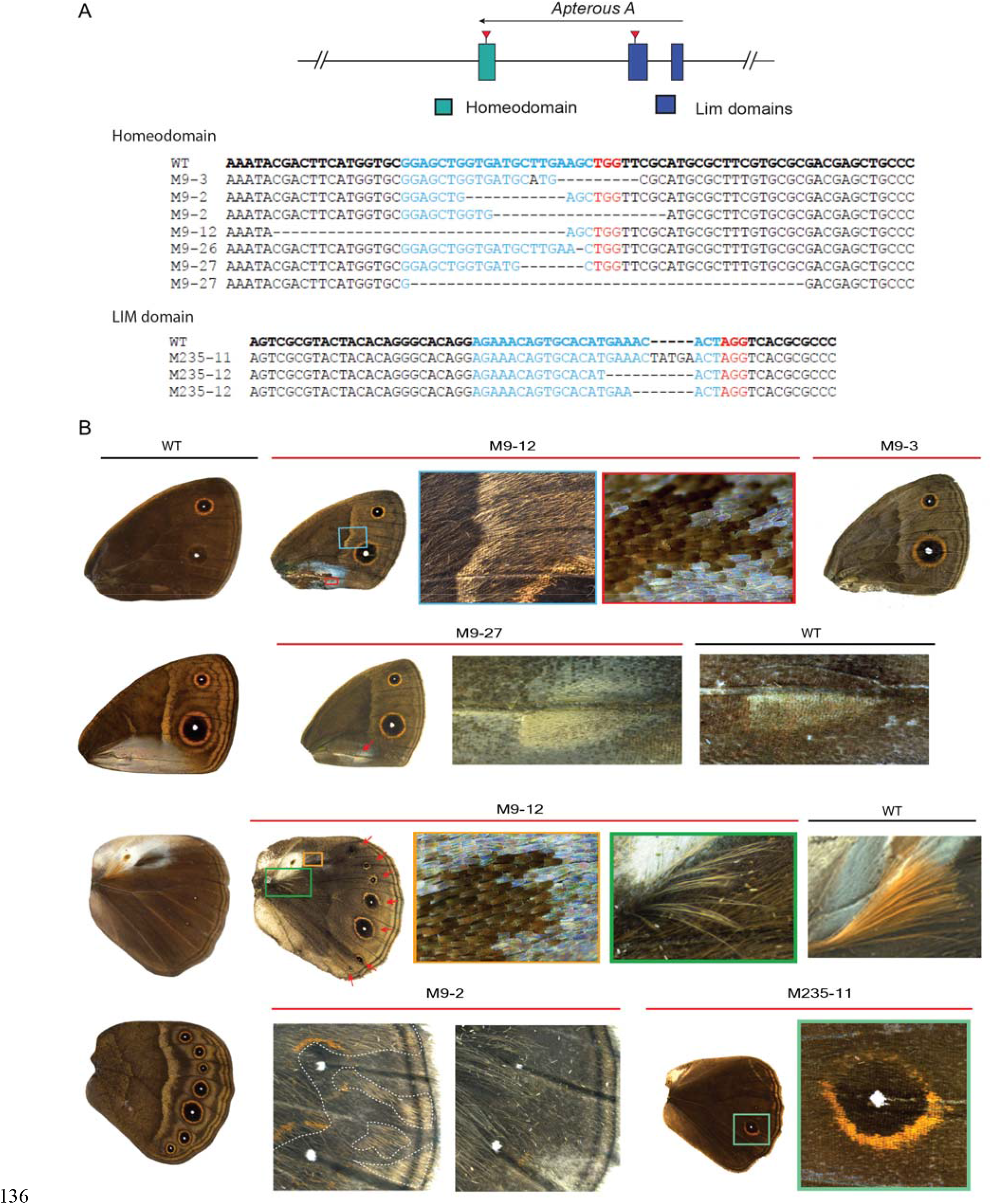
CRISPR/Cas9 mosaic wing pattern phenotypes of *apA* knockouts. A) Top: Regions of the *apA* gene in *B. anynana* targeted using the CRISPR/Cas9 system. Bottom: Sequences of the homeodomain and LIM domain regions of mutant individuals compared with the wildtype sequence in bold. Blue is the region targeted and the PAM sequence is in red. Deletions are indicated with ‘-‘. B) The range of CRISPR/Cas9 *apA* mutant phenotypes observed in *B. anynana*. The left column shows the wildtype (WT) dorsal and ventral surfaces for male forewings and hindwings. M9-12 (top): The dorsal forewing of a mutant male highlighting some of the ventral-like phenotypes and defects. The boxed regions are expanded to show the appearance of ventral-like white band and silver scales. M9-3: Dorsal forewing surface of a mutant female resembling the ventral surface. M9-27: Mutant with the ventral teardrop shape forewing androconial organ appearing on the dorsal surface (red arrow). WT dorsal forewing androconia is shown for comparison. M9-12 (bottom): A mutant dorsal hindwing with the appearance of all seven eyespots (red arrows), normally only seen on the ventral surface. The boxed regions are expanded to show the loss of silver scales associated with the dorsal hindwing androconia and improper development of hair-pencils. WT hair-pencil is shown for comparison. M9-2: Mosaic phenotype (left) on the dorsal surface with ventral-like light colored scales. Clones are indicated with a dashed white line. Corresponding region of the other wing of the same individual (right) shows no mosaicism. M235-11: A dorsal hindwing of a mutant with the width of the gold ring resembling that of ventral eyespots. Control animals, injected with only Cas9, all looked like wildtype (not shown).

No striking transformations of dorsal to ventral identity were observed in *apB* mutants. Some of the *apB* knockout phenotypes included wing hinge defect, a missing hindwing in one case (Fig S5: B-M9-22) and disturbed margin development (Fig S2: B-M9-17), sometimes associated with wing pattern disturbances (Fig S2: B-M9-15). Sequencing showed the presence of mutations in the targeted region (Fig S2A).

Knockdown of *apA* in a variety of insects from different lineages indicates that *apA* is necessary for wing growth and development and its function in this process seems to be highly conserved (5, 6, 8). However, our experiments, in agreement with others, also indicate a varying degree of co-option of this transcription factor into late wing development processes such as wing patterning and exoskeletalization. In *T. castaneum*, RNAi knockdown of *apA* and *apB* individually shows almost no phenotypic effects while their simultaneous knockdown leads to more dramatic phenotypes such as elytral exoskeletalization defects, depending on the developmental stage. Therefore, both *apA* and *apB* in beetles are important for early and late wing developmental processes (6). In *B. anynana*, knockout of both *apA* and *apB* causes defects in early wing development but only *apA* appears to have been co-opted to control dorsal surface-specific wing patterning.

### *apA* functions both as an activator and repressor of wing traits

Interestingly, our work shows that *apA* has multiple different, often antagonistic functions in surface- and sex-specific development between the fore- and hindwings. For example, *apA* acts as a repressor of male androconial organs and silver scale development on dorsal forewings, while it promotes hair-pencil and silver scale development on the dorsal hindwings of males (Fig 4A). These effects point to the likely interaction between *apA* and other factors such as sex-specific (*doublesex*) or wing-specific (*Ultrabithorax*) factors that together can specify sex- and surface-specific pattern development. We previously showed that *Ultrabithorax* (*Ubx)* is expressed in the hindwings but not forewings of *B. anynana* (9). In addition, the presence of a gene from the sex determination pathway, *doublesex* (*dsx*), in the future androconial regions of male wings of *B. anynana* was also verified by *in situ* hybridization and semi-quantitative PCR (10). These data support a likely combinatorial function reminiscent of the interactions between the hox gene *Scr* and *dsx* in the determination of the male-specific sex combs in the legs of *D. melanogaster* (11). The presence or absence of *Ubx*, type of *dsx* splice variant and *apA* may be sufficient to give each sex and wing surface a unique identity, though more work needs to be done to test this hypothesis. Given that proteins of the LIM-homeodomain subfamily, to which *ap* belongs, are unique in their ability to bind other proteins via their LIM domain (12), their involvement in such a large range of developmental processes, as repressors and activators, is likely.

**Figure 4:**
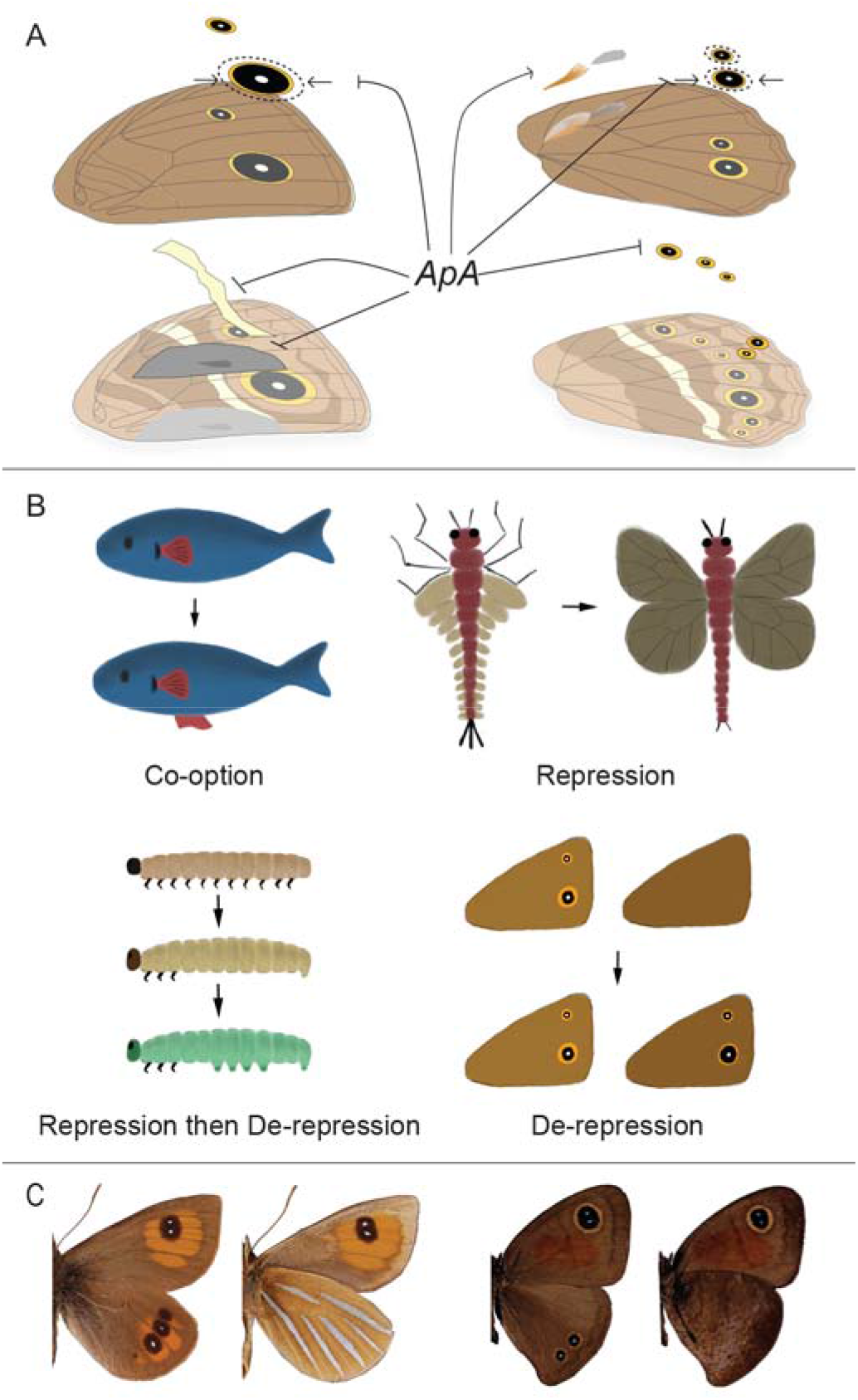
The role of *apterous* in surface-specific wing patterning in *B. anynana* and evolution of serial homologs in butterflies. A) A schematic of the different functions of *apA* on the dorsal surface of *B. anynana*. *apA* acts as a repressor of ventral traits such as the white transversal band, forewing androconia, hindwing eyespots, and the outer perimeter of the gold ring, and acts as an activator of hindwing hair-pencils and silver scales. B) Different modes of serial homolog evolution involving the co-option of a (fin) gene network to a novel body location (13), repression of the ancestrally repeated (wing) network in a subset of body segments (modified from (14)), repression followed by de-repression of the (limb) network in certain body segments (15), and de-repression of a never expressed (eyespot) network at a novel body location. C) *Argyrophenga antipodium* (left) and *Cassionympha cassius* (right) males with dorsal eyespots lacking ventral counterparts. Dorsal is to the left for each species.

### Discussion and Conclusion

Mutations in *apA* point to this gene functioning as a dorsal surface selector in *B. anynana* butterflies. Selector genes comprise a small set of developmental genes that are critical for specifying cell, tissue, segment, or organ identities in organisms (16).The wing selector hox gene *Ubx* allows hindwings to have a different identity from forewings. For example, the restricted expression of *Ubx* in hindwings of most insects examined so far, is required for membranous wing formation in beetles and bugs (17), haltere formation in flies (18) and hindwing specific color patterns in butterflies (19). When *Ubx* is mutated, in all the examples described above, hindwings acquire the identity of forewings, and when *Ubx* is over-expressed in forewings, these acquire a more hindwing-like identity (9). In *B. anynana*, *apA* functions in a similar manner along the dorsal-ventral axis of each wing – mutations in this gene make dorsal wing surfaces acquire a ventral identity. This type of homeotic mutation was also observed in a limited way, in bristles along the margin of the wings of *D. melanogaster*, where *ap* mutant clones developed bristles with a ventral identity (20). *B. anynana*, however, appears to have made inordinate use of *apA* for surface-specific color patterning and sexual trait development across the entire wing.

Further, this work highlights the possible role of *apA* in restricting the origin and early evolution of serial homologs such as eyespots in nymphalid butterflies to the ventral surface of the wings only. Broad comparative work across 400 genera of butterflies indicated that eyespots originated around 90 MYA within Nymphalidae on the ventral hindwing surface, and appeared ~40MY later on the dorsal surfaces (21–23). The appearance of additional eyespots on the dorsal surface of hindwings in *apA* mutants, and the absence of *apA* mRNA at the precise position where a few dorsal eyespots develop in both fore- and hindwings at the stage of eyespot center differentiation, implicates *apA* as a repressor of eyespot development in *B. anynana.* The additional gaps in *apA* expression observed in Spotty mutants further suggests that genetic mechanisms of eyespot number evolution on the dorsal surface proceeded via local repression of *apA*. We propose, thus, that the original ventral restriction of eyespots was due to the ancestral presence of *apA* on dorsal wing surfaces, and that eyespots’ later appearance on these surfaces was due to local *apA* repression.

The ancestral presence of a repressor (*apA*) of a gene regulatory network in a specific body location, followed by repression of the repressor, seems to represent a novel mode of serial homolog diversification (Fig 4B). This mode of serial homolog diversification is similar but also distinct from the mechanism previously proposed to lead to the re-appearance of abdominal appendages in lepidopteran larvae - via local repression of the limb repressor hox protein, *Abdominal-A* (*Abd-A*) (15, 24). In contrast to eyespots, when arthropod appendages first originated they were likely present in every segment of the body (25). Limbs were later repressed in abdominal segments, and finally they were de-repressed in some of these segments in some insect lineages (15). So, while the last steps of abdominal appendage and eyespot number diversification are similar (de-repression of a repressed limb/eyespot network), the early stages are different.

The comparative work across nymphalid butterflies also showed that the origin of dorsal eyespots was dependent on the presence of corresponding ventral eyespots in ancestral lineages (23). This implies that the extant diversity of eyespot patterns is biased/limited due to developmental constraints, probably imposed by *apA*. Interestingly, while ~99% of the species in our database display such constraints i.e dorsal eyespots always having ventral counterparts, a few butterflies display dorsal eyespots that lack ventral counterparts (Fig 4C). The molecular basis for these rare patterns remains to be explored.

In summary, we uncover a key transcription factor, *apA*, that due to its restricted expression on dorsal wing surfaces allowed *B. anynana* butterflies to develop and evolve their strikingly different dorsal and ventral wing patterns under natural and sexual selection. The interaction of *apA* with other sex- and wing-specific factors may explain the surface-specific pattern diversity we see across this as well as other butterfly species, but future comparative work is needed to further test these hypotheses. Additionally, our work has identified a new system to examine how developmental constraints, via *apA* repression of eyespot development, have shaped eyespot number biodiversity.

## Acknowledgments

We thank Arjen van’t Hof and Luqman Aslam for their help in retrieving sequence information from the *B. anynana* genome, Mainak Das Gupta for his help in the *in situ* hybridization protocols and Monteiro lab members for their support. This work was funded by Ministry of Education, Singapore grant R-154-000-602-112, the National University of Singapore grant R-154-000-587-133, and by the Department of Biological Sciences, NUS.

## Materials and Methods

### Animals

*Bicyclus anynana* butterflies were reared in a temperature controlled room at 27°C with a 12:12 hour light:dark cycle and 65% humidity. The larvae were fed on corn plants while the adults were fed on banana.

### Cloning and probe synthesis

*apA* sequence was obtained from [26] and *apB* and *dsx* sequences were identified from the *B. anynana* genome [27].The sequences were amplified with primers specified in Table S1, sequenced and then cloned into a PGEM-T Easy vector (Promega). Sense and anti-sense digoxigenin-labelled (DIG) riboprobes were synthesized *in vitro* using T7 and SP6 polymerases (Roche), purified by ethanol precipitation and resuspended in 1:1 volume of DEPC treated water:formamide.

### *In-situ* hybridization

The protocol was modified slightly from [28]. Briefly, larval or pupal wings were dissected from the last instar caterpillars or around 24-28 hrs after pupation respectively in PBS and transferred to glass well plates containing PBST (PBS+0.1% Tween20) at room temperature. The PBST was then immediately removed and the tissues fixed in 5% formaldehyde for 45 (larval) or 60 min (pupal) on ice, followed by 5 washes with cold PBST. The tissues were then incubated with 25µg/ml proteinase K in cold PBST for 4 (larval) or 5 minutes (pupal), washed twice with 2mg/ml glycine in cold PBST, followed by 5 washes with cold PBST. For larval wings, peripodial membrane was then removed on ice, post-fixed for 20 minutes with 5% formaldehyde and washed with PBST. The wings were gradually transferred to a prehybridization buffer (5X Saline sodium citrate pH 4.5, 50% formamide, 0.1% Tween20 and 100µg/ml denatured salmon sperm DNA), washed in the prehyb buffer and incubated at 60-65°C for 1 hour, followed by incubation in hybridization buffer (prehybridization buffer with 1g/L glycine and 70 to 140 ng/ml riboprobe) for 24 hours. The wings were then washed 6 to 10 times in prehybridization buffer at 60-65°C. They were then gradually transferred back to PBST at room temperature, washed 5 times in PBST and blocked overnight at 4°C (PBST+1% BSA). The DIG-labelled probes were then detected by incubating the tissues with 1:3000 Anti-DIG Alkaline Phosphatase (Roche) in block buffer for two hours, washed 10 times with block buffer, incubated in alkaline phosphatase buffer (100mM Tris pH 9.5, 100mM NaCl, 5mM MgCl_2_, 0.1% Tween) and finally stained with NBT/BCIP (Promega) solution at room temperature till colour developed. The reaction was stopped by washing in 2mM EDTA in PBST and again with PBST. The samples were either mounted on slides with ImmunoHistoMount medium (Abcam) or post-fixed with 5% formaldehyde before wax embedding and sectioning (Advanced Molecular Pathology Lab, IMCB, Singapore).

### Preparation of Cas9 mRNA and guide RNA

pT3TS-nCas9n was a gift from Wenbiao Chen (Addgene plasmid #46757). The plasmid was linearized with XbaI digestion and purified using a GeneJET PCR Purification Kit (Thermo Scientific). Cas9 mRNA was obtained by *in vitro* transcription using the mMESSAGE mMACHINE T3 kit (Ambion), tailed using the Poly(A) Tailing Kit (Ambion) and purified by lithium chloride precipitation. The guide RNA templates were prepared using a PCR based method according to [29]. The candidate targets were manually designed by searching for a GGN_18_NGG sequence on the sense or anti-sense strand of *apA* and *apB*, preferably targeting the LIM and homeobox domains of the transcription factor (Table S1). They were blasted against the *B. anynana* genome on LepBase.org to check for off-target effects. The template DNA sequence was used to perform an *in vitro* transcription using T7 RNA polymerase (Roche) at 37°C overnight, purified by ethanol precipitation and re-suspended in DEPC treated water.

### Microinjections

Eggs were collected on corn leaves within one to two hours of egg laying and were arranged on thin strips of double-sided tape on a petri dish. Cas9 mRNA and guide RNAs were mixed along with green food dye (Table S2) and injected into the eggs with a Borosil glass capillary (World Precision Instruments, 1B100F-3) using a Picospritzer II (Parker Hannifin). A piece of wet cotton was placed in the petri dish and the eggs were allowed to develop in an incubator at 27°C and high (~80%) humidity. Hatched caterpillars were placed on young corn plants using a brush. Adults that emerged were scored for their phenotypes (Table S2).

### Sequencing and genotyping mutants

Genomic DNA was extracted from leg tissues of mutant individuals using the E.Z.N.A Tissue DNA Kit (Omega Bio-tek). The region surrounding the target sequence was amplified by PCR, purified by ethanol precipitation, and used to check for presence of mutations using the T7 endonuclease I (T7EI) assay. Sequences from individuals with disruptions at the targeted regions were cloned into a PGEM-T Easy vector (Promega) and sequenced.

## Supplementary Materials

**Figure S1:**
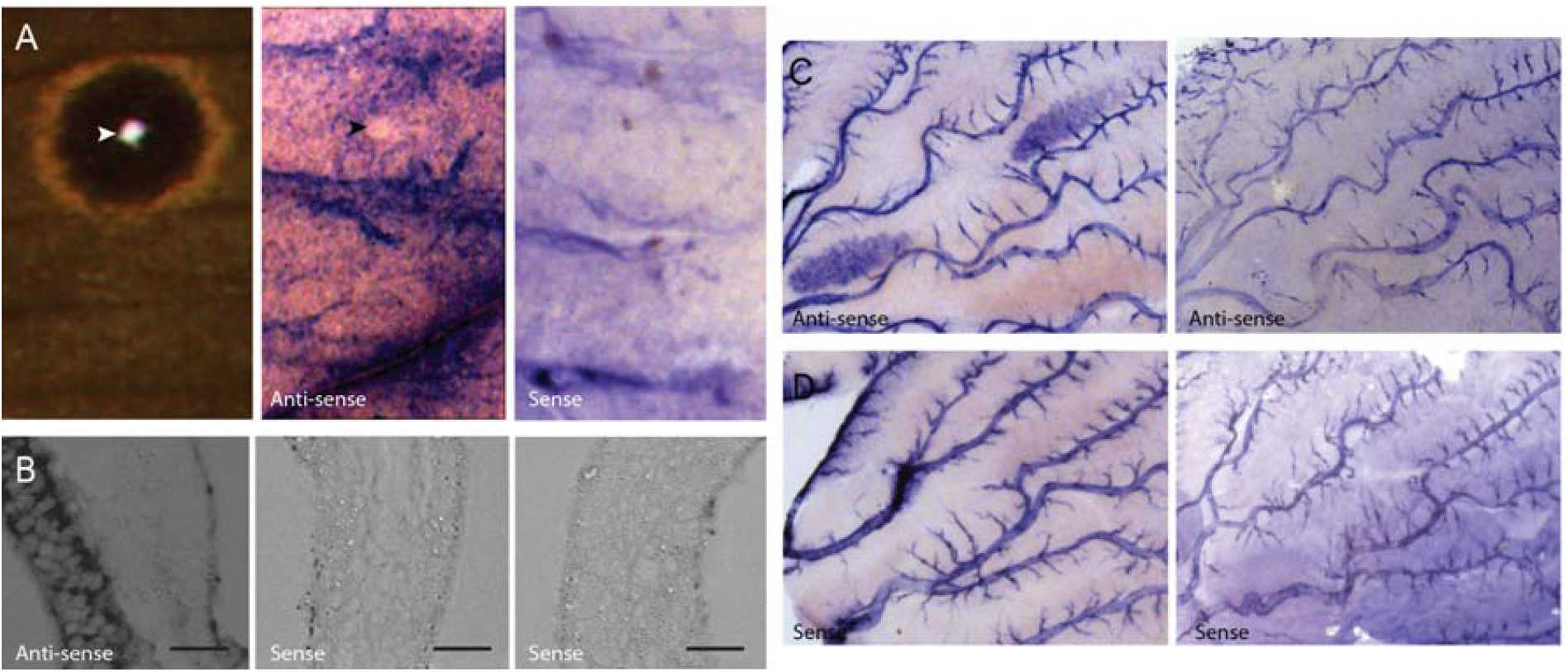
*ap* mRNA localization in developing wing discs of *Bicyclus anynana*. A) *apA* mRNA localization (middle) in wildtype 5^th^ larval instar wing discs with control (right). There is an absence of *apA* expression in future dorsal eyespot centers (arrowhead). Corresponding adult wing is shown (left). B) Cross-sectional view of a developing wing disc showing dorsal-specific *apB* expression (left). No staining is seen with control probes for *apB* (middle) and *apA* (right). Scale bar is 20µm C) Male (left) and female (right) hindwing discs (28 hours after pupation) showing *apB* mRNA up-regulation in the hair-pencil regions only in males. D) Controls for *apB* (left) and *apA* (right) expression in male wings show no staining in the corresponding regions.

**Figure S2:**
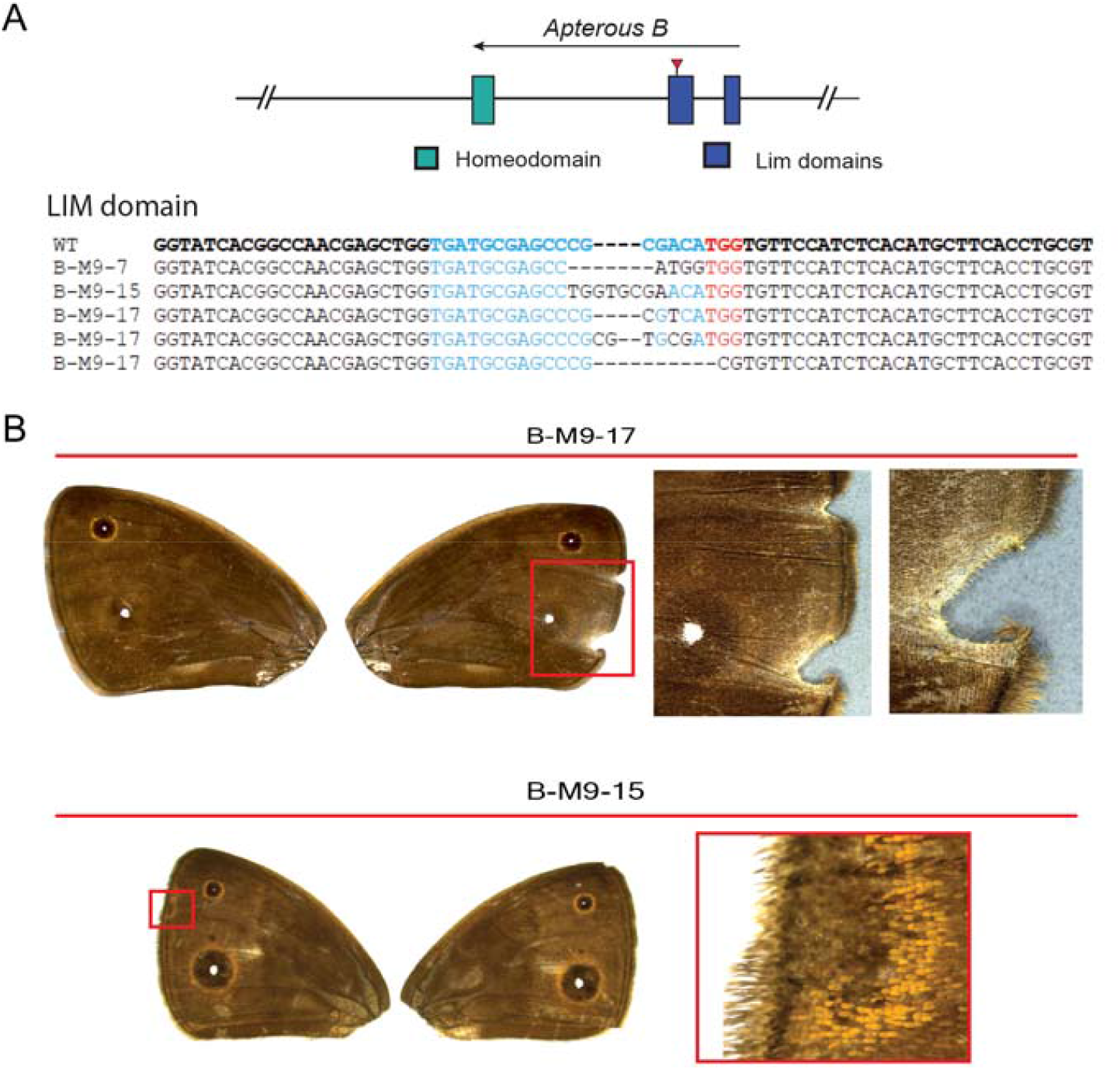
CRISPR/Cas9 mosaic wing pattern phenotypes of *apB* knockout. A) Top: Region of the *apB* gene in *B. anynana* targeted using the CRISPR/Cas9 system Bottom: Sequences of the LIM domain region of mutant individuals compared with the wildtype sequence in bold. Blue is the region targeted and the PAM sequence is in red. Deletions are indicated with ‘-‘. B) CRISPR/Cas9 *apB* mosaic phenotypes of *B. anynana*. B-M9-17: The forewings of a mutant individual showing differences in shape and marginal defects of the right wing as compared to the left. The boxed area is expanded to the right. B-M9-15: Mutant with wing pattern changes that do not correspond to mosaic ventral patterns, but appear to indicate disruptions to wing margin development. Boxed area expanded to the right.

**Figure S3:**
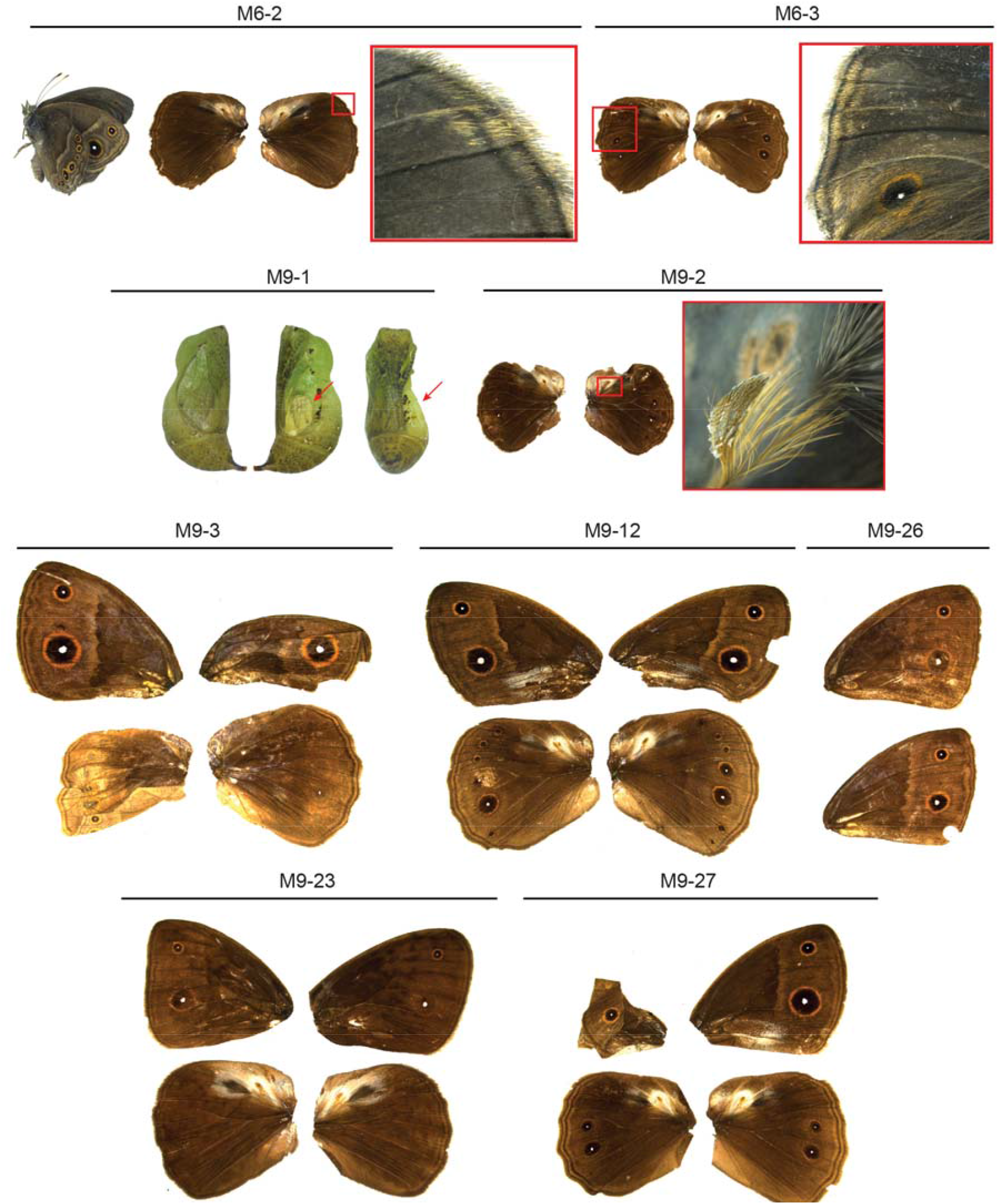
A catalog of all CRISPR/Cas9 mosaic wing pattern phenotypes of *apA* homeodomain knockout

**Figure S4:**
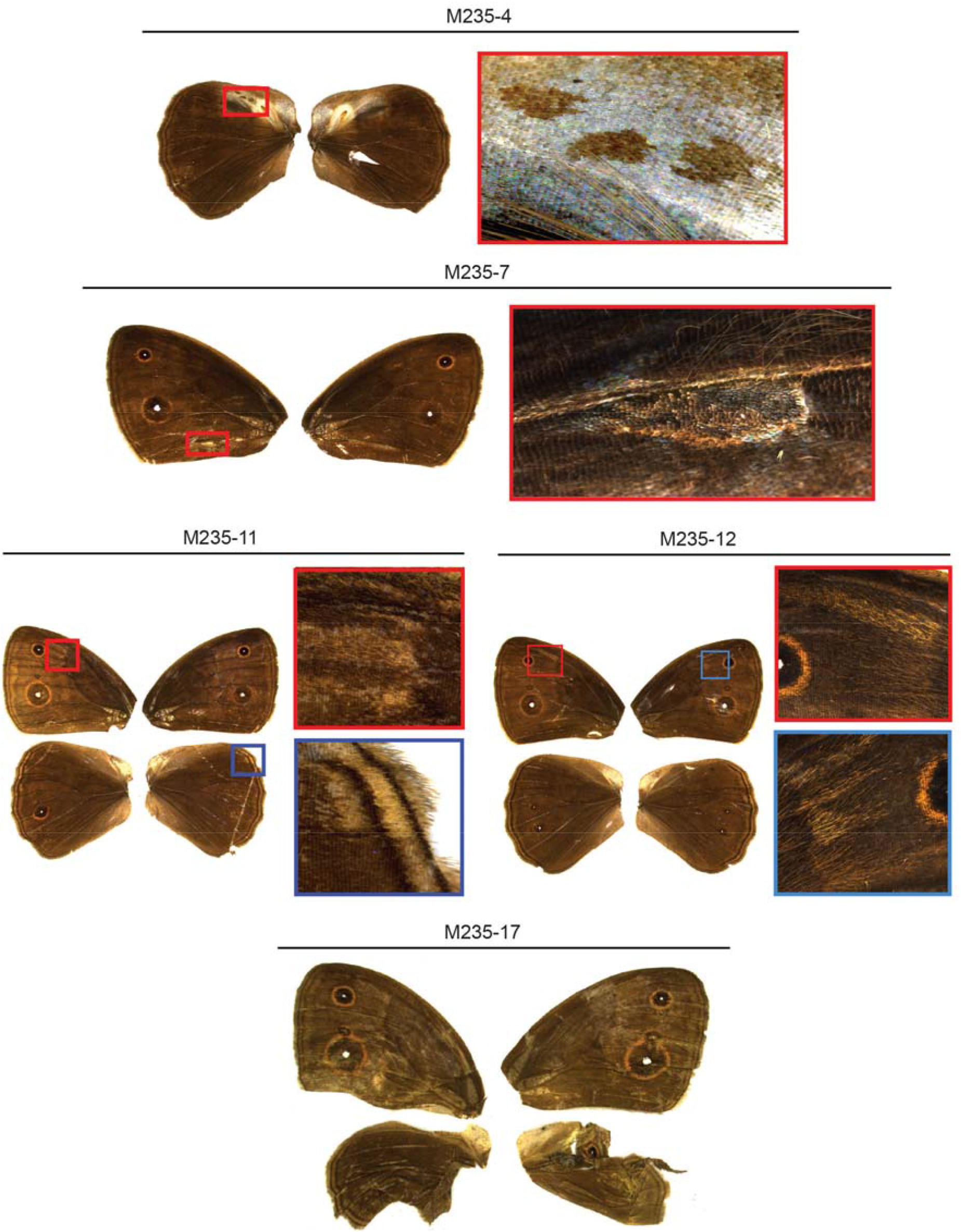
A catalog of all CRISPR/Cas9 mosaic wing pattern phenotypes of *apA* LIM domain knockout

**Figure S5:**
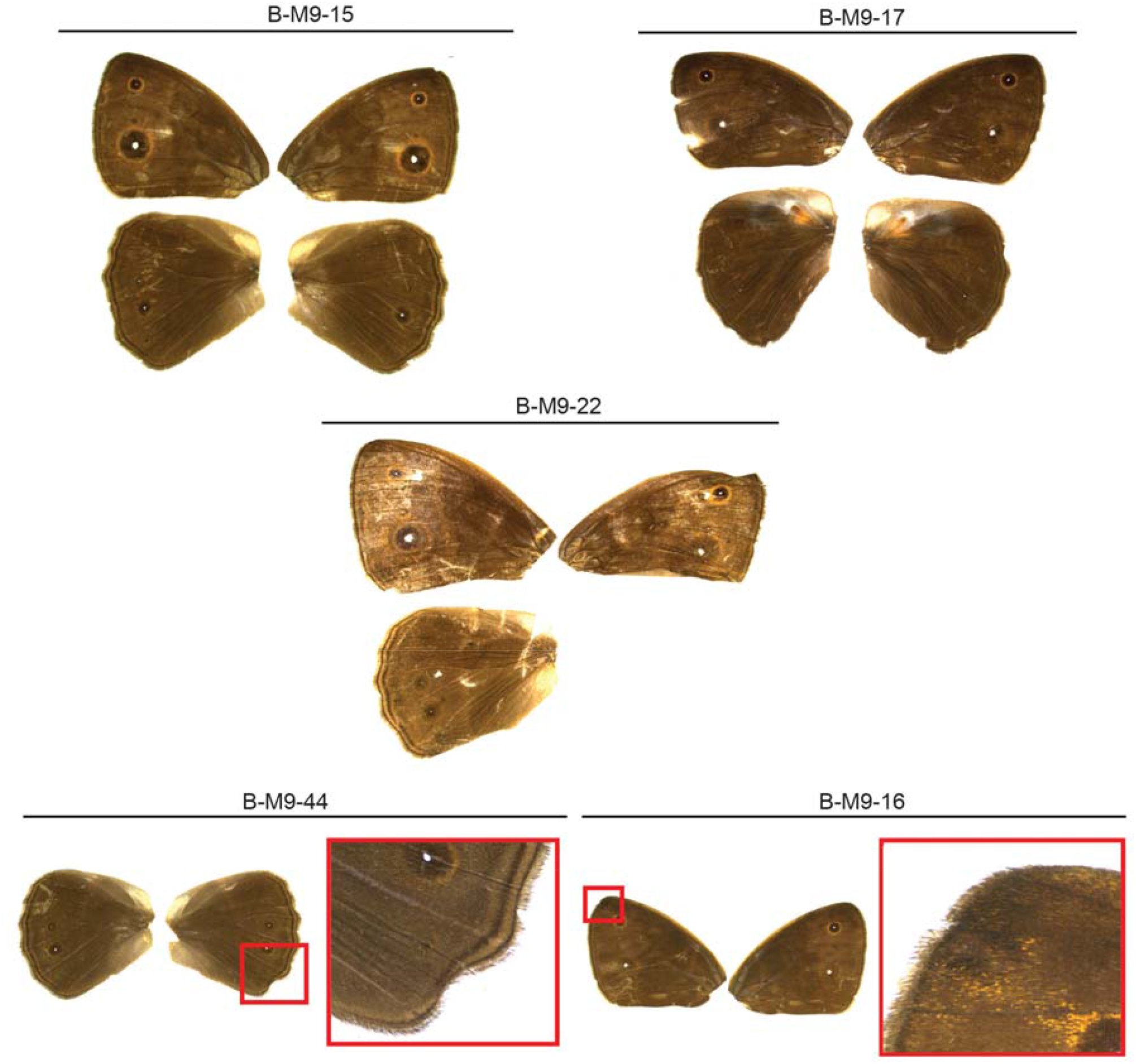
A catalog of all CRISPR/Cas9 mosaic wing pattern phenotypes of *apB* LIM domain knockout

**Table S1:**
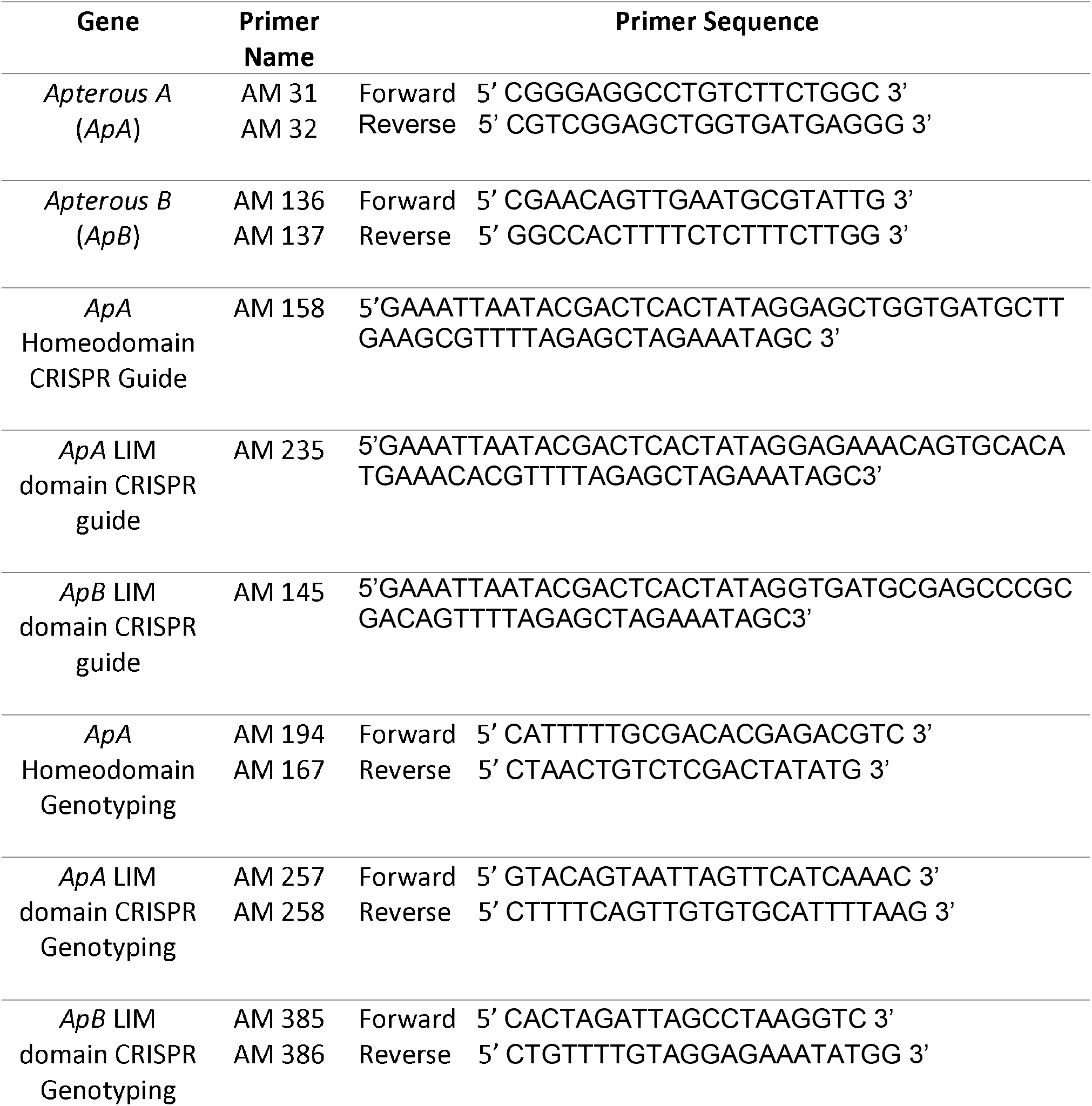
List of primers and guide RNA sequences used in this study

**Table S2:**
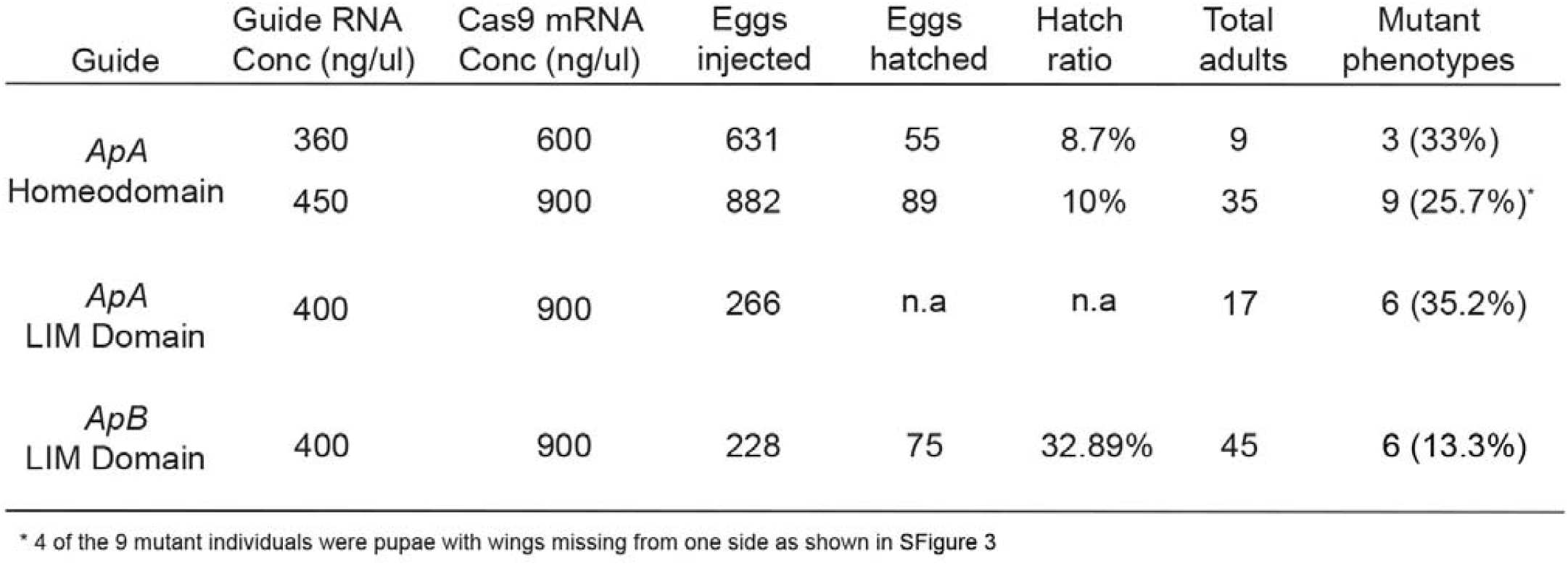
CRISPR/Cas9 injection concentrations and mutation frequencies

**Table S3:** CRISPR/Cas9 and control injection concentrations and hatch ratios

| Guide | Guide RNA<br>Conc (ng/ul) | Cas9 mRNA<br>Conc (ng/ul) | Eggs<br>injected | Eggs<br>hatched | Hatch<br>ratio |
| --- | --- | --- | --- | --- | --- |
| Control | - | 900 | 103 | 53 | 51.4% |
| *ApA*<br>Homeodomain | 400 | 900 | 113 | 75 | 66.3% |
| *ApA*<br>LIM Domain | 400 | 900 | 108 | 51 | 47.2% |
| *ApB*<br>LIM Domain | 400 | 900 | 104 | 53 | 50.9% |

